# Harnessing drought tolerance in a reference set of Andean amaranths

**DOI:** 10.1101/2025.08.06.668942

**Authors:** Ramiro. N. Curti, Jonatan Rodriguez, Pablo Ortega-Baes, Sergio J. Bramardi, Berta Velásquez, Alberto J. Andrade

## Abstract

Climate change and low-input farming systems increasingly expose crops to drought stress. Andean amaranths (*Amaranthus spp*.), as neglected and underutilized species, offer potential for climate-resilient agriculture due to their inherent drought tolerance and adaptability. In Northwest Argentina (NWA), a region with high environmental heterogeneity, exploiting this genetic diversity may improve food security. This study evaluated drought tolerance and yield stability among Andean amaranth genotypes to: assess the effects of genotype and G×E interaction, determine the potential for selecting specifically adapted genotypes, and identify high-yielding, stable genotypes for drought-prone conditions. Eleven genotypes (cultivars, breeding lines, and landraces of *A*. *caudatus* and *A*. *mantegazzianus*) were tested across four agroecological zones in NWA under irrigated and drought-stressed conditions. Grain yield data were analyzed using linear mixed models and AMMI analysis. Genotypes differed significantly in grain yield across environments and irrigation regimes. Strong G×E interactions led to genotype re-ranking across sites. Several *A*. *caudatus* breeding lines (G1, G2, G3, G6) combined high yield and stability. The *A*. *mantegazzianus* landrace (G11) was highly stable but low yielding. Amaranth genotypes showed distinct responses to drought, with some lines exhibiting broad adaptation and others, specific adaptation to stress-prone environments.

## Introduction

Many areas face critical food security challenges due to increasing agricultural demand and climate change’s impact on crop productivity (Lobell and Di Tommaso 2025). Among these impacts, drought stress stands out as a major abiotic stressor (Challinor *et al*. 2014). Significant advancements in breeding for crop yield in drought-prone regions have been achieved through extensive screening under abiotic stress (Ceccarelli 1994; Bänziger and Cooper 2001; Xangsayasane *et al*. 2014). Key strategies for breeding drought tolerance in low-input farming systems involve effective stress management, direct selection for specific adaptation, and the utilization of locally adapted germplasm (landraces) (Ceccarelli and Grando 1991; Bänzinger 2000; Ceccarelli *et al*. 2007). These principles suggest that selecting locally adapted germplasm can significantly improve breeding efficiency by exploiting genotype-by-environment (G×E) interactions (Hatfield and Walthall 2015). Furthermore, developing new varieties through participatory plant breeding is likely to translate into their fast adoption in low-input environments (Ceccarelli 2015).

Sustained use of genetic variability in neglected and underutilized species (NUS) offers a key adaptation strategy to mitigate effects of climate change (Padulosi *et al*. 2011). NUS are non-commodity crops marginalized for various reasons, yet they continuously evolve and adapt to environmental pressures (Andreotti *et al*. 2022). This ongoing adaptation makes them a dynamic source of alleles vital for crop improvement (Ulian *et al*. 2020). The Andean region, recognized as one of the world’s eight centers of domestication, boasts a high concentration of NUS (Galluzzi and López Noriega 2014). Among the various NUS seed crops, the pseudocereal group includes species from the genera *Chenopodium* and *Amaranthus* (family Amaranthaceae). These species are notable for their high nutritional quality and remarkable adaptability to a wide array of abiotic stresses (Joshi *et al*. 2018; Anuradha *et al*. 2023). Notably, amaranths exhibit drought resistance, requiring less water than many conventional staple crops (Pulvento *et al*. 2022; Gönen *et al*. 2024). This characteristic positions amaranths as a promising dryland crop for farmers in rainfed agricultural areas (Espitia-Rangel 1994; Pulvento *et al*. 2015).

The cultivated Andean amaranth grain includes the widely recognized species *Amaranthus caudatus*, alongside a taxonomically questioned entity known as *A*. *mantegazzianus*. First described by the Argentinean botanist Armando Hunziker in *Los pseudocereales de la agricultura indígena de America*, *A*. *mantegazzianus* was initially considered an endemic cultigen and the only domesticated species in Argentina. Its distribution is concentrated within the Calchaqui Valleys of Northwest Argentina, encompassing the provinces of Catamarca, Tucumán, and Salta (Hunziker 1952).

Despite later treatments suggesting *A*. *mantegazzianus* may be synonymous with *A*. *caudatus* or a derivative of *A*. *edulis* (Brenner *et al*. 2000; Alemayehu *et al*. 2015), its distinct panicle morphology sets it apart from other cultivated amaranths. Most cultivated amaranths exhibit a typical *amaranthiform* panicle, where glomeruli (flower clusters) are arranged along the secondary axis of the inflorescence. In contrast, *A*. *mantegazzianus* possesses a *glomerulate* panicle, characterized by glomeruli arranged on third-order axes. This unique panicle structure is also observed in quinoa (*Chenopodium quinoa*), another related pseudocereal originating from the Andean region (Curti *et al*. 2012). For the purpose of this study, we maintain the species identity of *A*. *mantegazzianus*, as early cytological studies validated it specific status (Greizerstein and Poggio 1994; Bonasora *et al*. 2013), and until further molecular studies evaluates its relationships within the Andean gene pool.

Field evaluations of amaranth germplasm have been conducted across various Latin American countries through the *Prueba Americana de Cultivares de Amaranto* to identify promising cultivars for high and stable seed yield (Mujica 1997). While these multi-environmental trials (METs) revealed a high potential for amaranth production, significant variability in crop yield was observed across locations, along with cultivar-specific responses (Mujica 1997). This highlights a substantial genotype-by-environment (G×E) interaction, complicating the selection of broadly adapted cultivars. This challenge is particularly pertinent in the Andean agricultural system, recognized as an extremely complex environmental mosaic (Jacobsen *et al*. 2003).

Here, diverse environmental factors, varying across toposequence and latitudinal range, profoundly influence crop yield and quality (Curti *et al*. 2014). Furthermore, Andean farmers typically operate with minimal inputs, cultivating amaranths on marginal lands where drought stress often represents a primary limiting factor for production (Kalinowski *et al*. 1992).

Northwest Argentina (NWA) is a vast region characterized by a diverse array of environments (Curti *et al*. 2012). To the west, the Cordillera Occidental, an extension of Bolivia’s Cordillera Real, introduces high-altitude landscapes known as the Puna. This ecoregion, while generally exhibiting consistently low year-round temperatures, is remarkably heterogeneous from north to south in terms of precipitation, temperature, and salinity (Ferrero and Villalba 2019). The Puna’s extremely short frost-free period severely limits the growing season, necessitating the cultivation of hardy crops capable of tolerating cold, drought, and saline conditions (Curti *et al*. 2012). Conversely, to the east, the Bolivian Cordillera Oriental extends into the region, creating high-altitude environments that are notably warmer than the Puna (Curti *et al*. 2012). Situated between these two mountain ranges, a series of gorges stretch from north to south, forming fertile areas that offer greater agricultural potential (Curti *et al*. 2012). During *Prueba Americana de Cultivares de Amaranto*, a test site in Purmamarca, Jujuy province, located in Northwest Argentina (NWA), demonstrated remarkable results. Despite a significant water deficit throughout the growing season, amaranth crops yielded acceptable grain (Mujica 1997). This observation suggests that the genetic variability in drought tolerance within Andean amaranth populations could be a valuable resource. This variability could be leveraged to develop new amaranth varieties specifically adapted to environments with varying water availability. Furthermore, given that NWA has experienced long-term shifts in precipitation patterns, characterized by changes in both frequency and amount (Ferrero and Villalba 2019), the ongoing search for amaranth germplasm with desirable levels of drought tolerance is highly encouraged.

This study aims to screen genotypic variability for drought stress tolerance among Andean amaranth cultivars, breeding lines, and landraces within managed stress environments. We hypothesize that by managing drought stress through irrigation differences, we will observe the largest genotypic variation, and productivity will be maximized by selecting genotypes for specific environments. Specifically, we addressed the following objectives: i) to examine the relative magnitude of genotype (G) and genotype by environment (G×E) interaction effects both between and within Andean amaranth species; ii) to evaluate the relative advantage of selecting for specific adaptation by exploiting G×E interaction; and iii) to identify high-yielding and stable Andean amaranth genotypes suitable for drought-prone environments.

## Material and Methods

### Genotypes and testing locations

A reference set of eleven amaranth genotypes, comprising six breeding lines, one landrace and four cultivars of *Amaranthus caudatus* and one landrace of *A*. *mantegazzianus* (Table 1), were evaluated in four sites in the province of Jujuy (Argentina). Genotype selection was based on diversity of origin and most came from the *Prueba Americana de Cultivares de Amaranto*, an international METs in which a set of genetically diverse cultivars was tested in several countries of Latin America (Mujica 1997). The genotypes originated from different breeding programs (Table 1), and a short description of their pedigree follows. Breeding lines 10c, 294ab, 32a, 38a, 44a and G403 and cultivar Noel Vietmayer were selected from ecotypes collected in Tarija (Bolivia) at Centro de Investigaciones en Cultivos Andinos (CICA), Cuzco (Peru) (Kalinowski *et al*. 1992). Cultivar INIAP-alegría was selected from a local landrace at Instituto Nacional de Investigaciones Agropecuarias (INIAP, Quito, Ecuador) for cultivation at mid-elevation valleys (Nieto 1993). Cultivar UTAB- Cahuayuma was selected from a local landrace at Estación Experimental Pairumani, Instituto Boliviano de Tecnología Agropecuaria (IBTA; currently PROINPA) (Rojas *et al*. 2010). The CVR genotype derived from mass selection of a local landrace collected in Jujuy (Argentina), whereas genotype MTGZ correspond to another amaranth seed crop (*A*. *mantegazzianus*) which was selected from a local landrace collected in Jujuy (Argentina) (Table 1).

**Table 1.** Code, species, origin and status of the 11 genotypes of Andean amaranths evaluated in multi-environment trials in Northwest Argentina.

| Code | Species | Genotypes | Origin | Status |
| --- | --- | --- | --- | --- |
| G1 | *A. caudatus* | 10c | Cuzco, Perú | Breeding line |
| G2 | *A. caudatus* | 294ab | Cuzco, Perú | Breeding line |
| G3 | *A. caudatus* | 32a | Cuzco, Perú | Breeding line |
| G4 | *A. caudatus* | 38a | Cuzco, Perú | Breeding line |
| G5 | *A. caudatus* | 44a | Cuzco, Perú | Breeding line |
| G6 | *A. caudatus* | G403 | Cuzco, Perú | Breeding line |
| G7 | *A. caudatus* | CVR | Jujuy,<br>Argentina | Landrace |
| G8 | *A. caudatus* | INIAP-alegría | Ecuador | Cultivar |
| G9 | *A. caudatus* | NOEL-Vietmeyer | Perú | Cultivar |
| G10 | *A. caudatus* | UTAB-Cahuayuma | Bolivia | Cultivar |
| G11 | *A. mantegazzianus* | MTGZ | Jujuy,<br>Argentina | Landrace |

The experimental sites were located in farmer’s fields including the major agroecological zones in which amaranth crops grown in the province of Jujuy. Accordingly, the sites represent the Target Population of Environments (TPEs) into which the amaranth breeding program operates. The León site (Department of Manuel Belgrano), located at low elevation (1754 m asl) represents a typical humid valley environment. Purmamarca (Department of Tumbaya) and Hornaditas (Department of Humahuaca) sites represents a typical dry valley environment. Purmamarca is located at a higher elevation than León (2324 m asl) but at lower than Hornaditas (3230 m asl). The last site selected, La Quiaca (Department of Cochinoca), is located at high elevation (3440 m asl) and represents a typical Puna environment. Precipitation and evapotranspiration values for the entire growing season were obtained from meteorological stations located at 500 m distance from the experimental sites.

### Field trials

Two trials were planted side by side at each experimental site at a safe distance of 40 to 50 m. Between trials, corn (*Zea mays* L.) was sown to avoid the neighbor effect of the treated trial on the untreated trial. In each of the two trials, all genotypes were planted in the field in a randomized complete block design with four replicates. The first trial served as a control and received deep irrigation before planting and then every 10 or 15 days following local practices (furrow irrigation) to avoid water deficits during the crop cycle. The second trial was drought stressed by stopping irrigation between the 85 and 125 days after crop emergence at the stage of anthesis (i.e., when at least one flower is open, corresponding to stage R4 in the phenological scale of Henderson *et al*. (1998). After the stage of the end of anthesis the irrigation was reinitiated in the drought stress trial in the same fashion as the control trial. Plots were three rows with 70 cm inter-row spacing and 5 m long. Sowing density was 47,620 grains ha^-1^ and weeds were removed by hand and fungicides and insecticides applied upon diseases and pest detection in the field. The experiments were fertilized at a rate of 43 kg N ha^-1^ after crop emergence. At physiological maturity (visually determined by the change in panicle color, corresponding to stage R7 in the phenological scale of Henderson *et al*. (1998)), ten contiguous plants from each replicate were harvested to record grain yield adjusted to 14% of moisture content and expressed in tones ha^-1^.

### Statistical analysis

A linear mixed model was set up for an analysis of grain yield across trials and sites as follow:

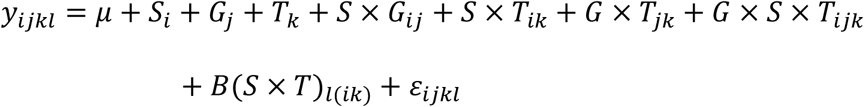

with *y* designating the grain yield of the *j-*th genotype in *i-*th site, *k-*th trial and *l-*th block, 𝜇 is the grand mean, *S* is the effect of *i-*th site, *G* is the effect of *j-*th genotypes, *T* is the effect of *k-*th trial (drought vs. irrigation), *GS_ij_* is the interaction effect of *j*-th genotype and *i*-th site, *ST_ik_* is the interaction effect of *i*-th site and *k*-th trial, *GT_jk_* is the interaction effect of the *j*-th genotype and *k*-th trial, *GST_ijk_* is the triple interaction effect of genotype *j*-th and *i*-th site and *k*-th trial, *B* the random effect of *l*-th block nested within the *i*-th site by *k*-th trial, and 𝜀 the random error. In the model, the genotypes, trials and sites were considered fixed whereas blocks were considered random. The analysis was computed using the PROC GLM in SAS® software (SAS Institute 2004).

In a second-stage, we restructured the G×S×T dataset into a two-way G×E dataset, treating each unique combination of location and trial as a distinct environment. We then performed an Additive Main and Multiplicative Interaction (AMMI) analysis using the metan package (Olivoto and Lúcio 2020) within the R statistical environment (R core team, 2024). This restructuring was necessary because while the ANOVA component of AMMI can handle factorial designs of any complexity, the principal components analysis (PCA) part of the AMMI model specifically requires a two-way dataset (Paderewski *et al*. 2016). The aim was to model the genotype-by-environment interaction to describe the pattern of genotypic response and identify high yielding and stable genotypes. The number of IPCA (Interaction Principal Components Analysis) axes to retain in the AMMI model was based on the Gollob’s test. The Biplots were constructed for the AMMI1 and AMMI2 models. We computed the Weighted Average of Absolute Scores (WAAS, Olivoto *et al*. (2019)) to determine the stability of genotypes and the discriminant ability of sites. Finally, with the computed WAASBY index to get insight into the simultaneous selection for mean performance and stability (Olivoto *et al*. 2019).

## Results

In the León site, precipitation exceeds evaporation for the majority of the growing season, resulting in negligible water deficit, with the exception of the late flowering stage (Fig. 1a). In contrast, the Purmamarca and Hornaditas sites exhibit evaporation values exceeding precipitation throughout the entire growing season, leading to substantial water deficits, particularly during the flowering period (Fig. 1b and c). At La Quiaca, precipitation exceeds evaporation only at the start of the growing season, while a high-water deficit was observed during the flowering period as evaporation increases towards the season’s end (Fig. 1d).

**Fig. 1.**
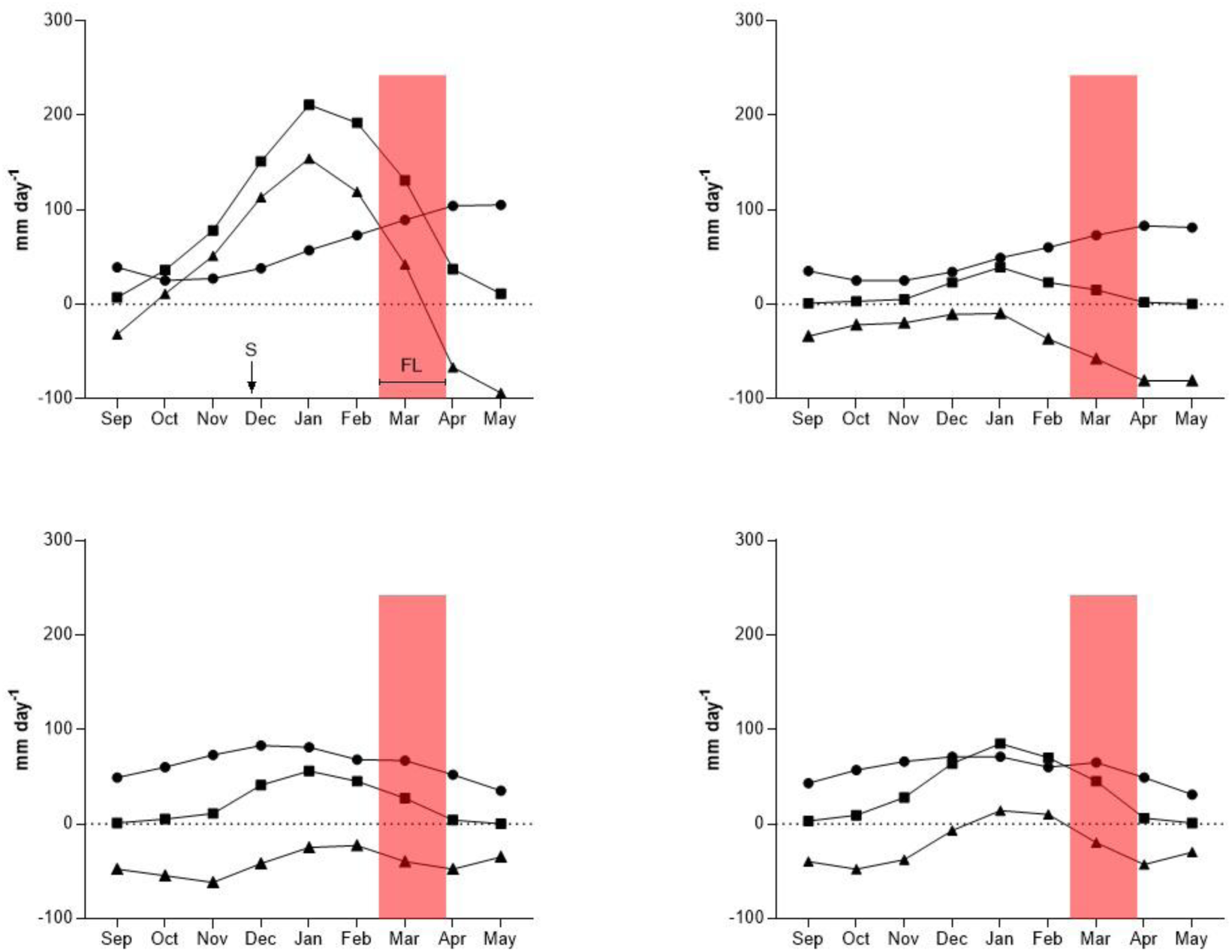
Mean rates of precipitation (P) and evaporation (E) and the difference between them (P-E) (mm water day^-1^) for the four sites in which Andean amaranths were evaluated: (a) León; (b) Purmamarca; (c) Hornaditas and (d) La Quiaca. The sowing time (S) and the flowering period (FL) are shown.

A statistically significant genotype-by-location-by-trial interaction effect was observed for grain yield among the Andean amaranth genotypes evaluated in Northwest Argentina (Table 2). The performance plots illustrated a consistent re-ranking of genotypes across different sites between irrigated and drought-stress trials, indicating strong crossover interactions (Figure S1). The Additive Main effects and Multiplicative Interaction (AMMI) model corroborated the G×E interaction effect and revealed that the first five Interaction Principal Component Axes (IPCA) were significant in explaining the variation in genotype performance across environments (Table S2).

**Table 2.** The ANOVA for genotype × site × trial adjusted means of Andean amaranth grain yield.

| Sources of variation | df | Sum of squares | Mean squares | *F*-ratio | *P* |
| --- | --- | --- | --- | --- | --- |
| Trial (T) | 1 | 9.34 | 9.34 | 400.81 | <.0001 |
| Site (S) | 3 | 2.43 | 0.81 | 34.84 | <.0001 |
| S×T | 3 | 3.5 | 1.16 | 50.07 | <.0001 |
| Genotype (G) | 10 | 10.33 | 1.03 | 44.33 | <.0001 |
| G×T | 10 | 0.59 | 0.05 | 2.55 | 0.0061 |
| T×B (S) | 12 | 0.32 | 0.02 | 1.17 | 0.308 |
| G×S | 30 | 2.48 | 0.08 | 3.56 | <.0001 |
| G×S×T | 30 | 3.01 | 0.1 | 4.32 | <.0001 |

In the AMMI1 model, environments E4 and E6 were positioned on opposing sides along the IPCA axis, demonstrating a pronounced influence on their capacity to differentiate among genotypes (Fig. 3a). The remaining environments exhibited short vector lengths, implying a limited discriminatory power along the first IPCA. Genotypes G2, G3, and G11 were located near the origin of the IPCA axis, suggesting their yield stability across all evaluated environments (Fig. 3a). Notably, G2 displayed the highest grain yield values, while G11 exhibited the lowest. Genotypes G1, G4, G5, G6, G7, G8, and G9 demonstrated cross-over interaction between E4 and E6, indicating specific adaptation to these environments (Fig. 2a). Genotypes G1, G5, and G10 showed high grain yield than grand mean, whereas G9 showed low grain yield, however, all exhibited improved performance in environment E4 (Fig. 2a).

Conversely, genotypes G6 and G7, which had high yields, and G4 and G8, which had low yields, showed improved performance in environment E6 (Fig. 2a).

**Fig. 2.**
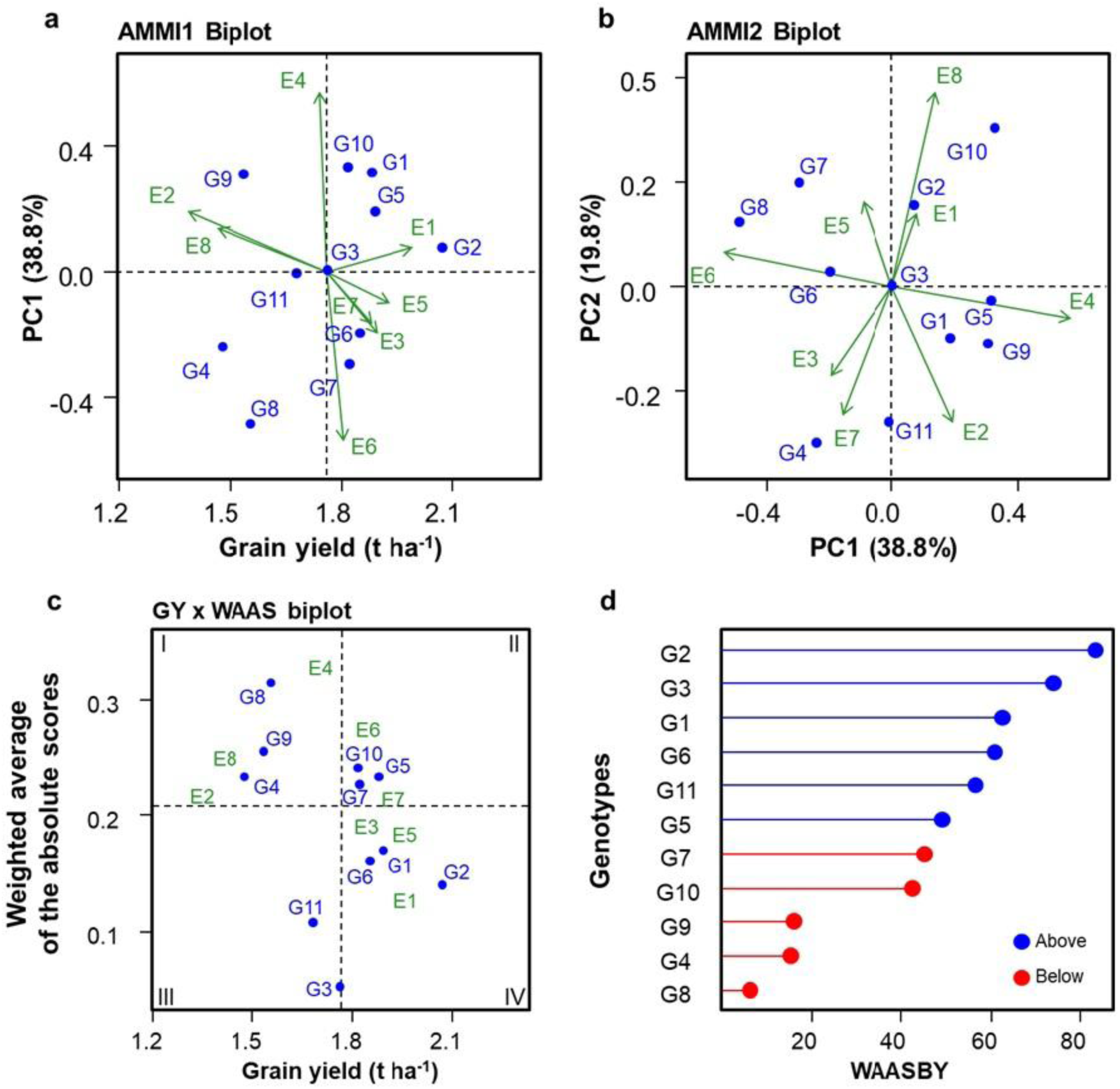
Additive Main effects and Multiplicative Interaction (AMMI) model for amaranth grain yield and stability analysis. AMMI for the: (a) first IPCA plotted against grain yield (AMMI1) and, (b) the first two IPCA2 (AMMI2); (c) stability analysis based on Weighted Average of Absolute Scores against grain yield; and (d) ranking of genotypes based on WAASBY index and the opportunity to simultaneous selection for mean performance and stability. Genotypes (G#) are denoted by symbols (circle) and Environments (E#) by vectors, E1: León irrigated; E2: León stressed; E3: Purmamarca irrigated; E4: Purmamarca stressed; E5: Hornaditas irrigated; E6: Hornaditas stressed; E7: La Quiaca irrigated; E8: La Quiaca stressed.

In the AMMI2 model, the environments spanned the entire Euclidean space, underscoring the substantial G×E interaction effects. Consistent with the AMMI1 model, environments E4 and E6 were situated on opposing sides along the IPCA1 axis, demonstrating the same contrasting influences (Fig. 2b). Furthermore, the IPCA2 axis revealed contrasting effects for environments E8 and E7. Environments E3 and E2 exhibited a close relationship with E7, as indicated by angles less than 90° (Fig. 2b).

Conversely, E1 and E5 displayed short vectors, suggesting that their capacity to capture G×E interactions would likely be accounted for by higher-order IPCAs (Fig. 2b). Genotypes G1, G5, and G9 showed improved performance in environment E4, while G6, G7, and G8 performed better in E6. Conversely, G4 and G11 exhibited improved performance towards E2, E3, and E7, whereas G2 and G10 showed improved performance towards E8 (Fig. 2b). Notably, G3 was the only genotype exhibiting stability, as it was positioned at the origin of both the IPCA1 and IPCA2 axes (Fig. 2b).

The biplot of Weighted Average of Absolute Scores (WAASB) values plotted against mean grain yield facilitates a four-quadrant classification system for the concurrent interpretation of yield performance and stability (Fig. 2c). Genotypes G4, G8, and G9, along with environments E2, E4, and E8, situated in quadrant I, are characterized as unstable genotypes or environments exhibiting high discriminatory power but with productivity below the overall mean (Fig. 2c). Similarly, genotypes G5, G7, and G10, and environments E6 and E7, located in quadrant II, also demonstrate instability and high discrimination ability, yet their productivity surpasses the grand mean (Fig. 2c). Genotype G11, positioned in quadrant III, displays low productivity but is considered stable due to its lower WAASB values (Fig. 2c).

Conversely, genotypes G1, G2, G3, and G6, and environments E1, E3, and E5, found in quadrant IV, are identified as highly productive and broadly adapted, owing to their high mean yield and superior stability performance (lower WAASB values) (Fig. 2c). The genotype ranking considering both the stability (WAAS) and mean performance based on the WAASBY index confirmed the high productive and broadly adaptation of genotypes G1, G2, G3 and G6 (Fig. 2d).

## Discussion

We applied managed drought stress through irrigation at the flowering stage and found that it differentially affected the performance of Andean amaranth genotypes across sites. This revealed clear genotypic variation in drought tolerance. Sensitivity to stress also varied with the climatic conditions at each site. Most genotype-by-environment (G×E) interactions resulted in a reranking of genotype performance, complicating the development of effective selection strategies. Therefore, improving grain yield in Andean amaranths requires breeding programs to explicitly address the G×E interactions observed within Target Population of Environments (TPEs) from Northwest Argentina.

Drought sensitivity around flowering in Andean amaranths is comparable to other grain or vegetable amaranth species cultivated in North America and Asia (Johnson and Henderson 2002; Jamalluddin *et al*. 2021). Furthermore, yield reductions under managed stress highlight that flowering stage as critical for yield determination (Gönen *et al*. 2024). The study also found performance differences between species, especially within *A*. *caudatus*, which showed significant genetic variability in drought response. Future studies in amaranth research should focus on understanding the morpho-physiological mechanisms that explain varying sensitivities to drought stress within this species. Prior research on other *Amaranthus* species offers valuable insights. For instance, *A*. *cruentus* and *A*. *hypochondriacus* have demonstrated enhanced root exploration under drought conditions, leading to improved water use efficiency (Johnson and Henderson 2002). In contrast, *A*. *hybridus* and other vegetable amaranth varieties employ a different strategy, reducing stomatal conductance as a water conservation mechanism when exposed to drought and salinity stresses (Liu and Stützel 2002, 2004; Omamt *et al*. 2006; Jamalluddin *et al*. 2019). Therefore, future field experiments should prioritize investigating root exploration in Andean amaranth species under drought stress. This research will be crucial for developing more drought-resilient amaranth varieties, which is vital for food security in regions prone to water scarcity.

Our findings reveal that *A*. *caudatus* genotypes at different breeding stages exhibit distinct performance patterns, indicating varied adaptive strategies across the TPEs. The first Interaction Principal Component Axis (IPCA1) of the AMMI1 model demonstrated that G2 and G3, both *A*. *caudatus* breeding lines (Table 1), displayed contrasting responses. G3 showed high stability across environments, while G2 exhibited higher grain yield (Fig. 2a). Conversely, G11, a landrace of another amaranth species (Table 1), was positioned near the origin, signifying consistent performance across environments but lower grain yield. This aligns with previous research indicating that *A*. *mantegazzianus* can achieve acceptable yields in Northwest Argentina with approximately 176 mm of accumulated rainfall (Mujica 1997). The current study’s environments had rainfall ranging from 111 to 190 mm (Fig. 1b and c), further supporting the adaptability of *A*. *mantegazzianus* under such conditions.

The AMMI2 biplot confirmed the high stability of breeding line G3. In general, the *A*. *caudatus* cultivars displayed close adaptation to specific environments along the first IPCA1. This is exemplified by the contrasting responses of G8, G9, and G10 between E4 and E6 (Fig. 2b), which were the most drought-stressed environments evaluated (Fig. 1b and c). A similar performance pattern was observed for other *A*. *caudatus* breeding lines (G1, G5, and G6) and the landrace G7 (Fig. 2b). On the other hand, G2 and G11 exhibited divergent environmental responses along the second IPCA. Specifically, G2 demonstrated improved performance in environments E1, E5, and E8, whereas G11 in E2, E3, and E7 (Fig. 2b). These environmental groupings appear geographically and climatically disparate, as they varied in altitude, accumulated precipitation, and management practices. Consequently, despite observing strong crossover G×E interactions, the results do not support their subdivision into distinct mega-environments. These findings diverge from those reported for quinoa, in which evaluations conducted in analogous environments revealed that the delineation into mega-environments remained consistent, irrespective of alterations in management practices (Curti *et al*. 2014; Agüero *et al*. 2023).

Our results reveal a clear distinction between environments based on their discriminatory capacity and average grain yield. Environments characterized by high discriminatory capacity but lower average yields were predominantly drought-stressed (E2, E4, and E8). Conversely, irrigated environments (E6 and E8) exhibited higher average grain yields. All evaluated cultivars, the landrace, and some breeding lines of *A*. *caudatus* were associated with these high-yielding, irrigated environments, yet they displayed greater instability. Notably, no environment fell into Quadrant III (low discriminatory capacity and low grain yield) in our analysis (Fig. 2c). Interestingly, G11 demonstrated superior stability (Fig. 2c). Conversely, irrigated environments (E1, E3, and E5) showed low discriminatory capacity but higher-than-average grain yields. The remaining *A*. *caudatus* breeding lines were associated with these stable environments, exhibiting consistent performance across the target population of environments (TPEs). A comprehensive evaluation of genotypes based on their average grain yield and stability clearly indicated the higher stability of *A*. *caudatus* breeding lines and the *A*. *mantegazzianus* landrace, while *A*. *caudatus* cultivars demonstrated specific adaptation (Fig. 2d).

Future research should explore the potential of exploiting genotype-environment (G×E) interactions for specific adaptation within the Northwest Argentina TPEs. This could involve focusing on the observed ranking changes in *A*. *caudatus* cultivars and its landrace, and, to some extent, selecting breeding lines that exhibit crossover interactions. Alternatively, a breeding strategy that minimizes G×E interaction and prioritizes the selection of genotypes for wide adaptation and high stability remains a viable option. Subsequent studies should assess the logistical and resource demands of implementing each of these breeding strategies within these complex TPEs.

## Data Availability Statement

The data that support this study will be shared upon reasonable request to the corresponding author.

## Supporting information

Plot of performance for Andean amaranth genotypes evaluated in paired trials under irrigation and stressed conditions

The ANOVA for the additive main effects and multiplicative interaction model

## Acknowledgments

We would like to extend our thanks to all the farmers involved in the evaluation of amaranth materials.

## References

Agüero JJ, Acreche MM, Sühring SS, Bertero HD, Curti RN (2023) Genotype-dependent responses of Andean and Coastal quinoa to plant population density for yield and its physiological determinants in Northwest Argentina. Crop & Pasture Science 75, CP23040. 10.1071/CP23040

Alemayehu FR, Bendevis MA, Jacobsen S -E. (2015) The potential for utilizing the seed crop amaranth (*Amaranthus* spp.) in East Africa as an alternative crop to support food Security and climate change mitigation. Journal of Agronomy and Crop Science 201, 321–329. 10.1111/jac.12108

Andreotti F, Bazile D, Biaggi C, Callo-Concha D, Jacquet J, Jemal OM, King OI, Mbosso C, Padulosi S, Speelman EN, van Noordwijk M (2022) When neglected species gain global interest: Lessons learned from quinoa’s boom and bust for teff and minor millet. Global Food Security 32, 100613. 10.1016/j.gfs.2022.100613

Anuradha, Kumari M, Zinta G, Chauhan R, Kumar A, Singh S, Singh S (2023) Genetic resources and breeding approaches for improvement of amaranth (*Amaranthus* spp.) and quinoa (*Chenopodium quinoa*). Frontiers in Nutrition 10. 10.3389/fnut.2023.1129723

Bänziger M, Cooper M (2001) Breeding for low input conditions and consequences for participatory plant breeding examples from tropical maize and wheat. Euphytica 122, 503–519.

Bänzinger M, Edmeades GO, Beck DL, Bellon MR (2000) Breeding for drought and nitrogen stress tolerance in maize: From theory to practice. CIMMYT

Bonasora MG, Poggio L, Greizerstein EJ (2013) Cytogenetic studies in four cultivated *Amaranthus* (Amaranthaceae) species. Comparative Cytogenetics 7, 53–61. 10.3897/CompCytogen.v7i1.4276

Brenner DM, Baltensperger DD, Kulakow PA, Lehmann JW, Myers RL, Slabbert MM, Sleugh BB (2000) Genetic resources and breeding of *Amaranthus*. Plant Breeding Reviews 19, 227–285.

Ceccarelli S (1994) Specific adaptation and breeding for marginal conditions. Euphytica 77, 205–219.

Ceccarelli S (2015) Efficiency of plant breeding. Crop Science 55, 87–97. 10.2135/cropsci2014.02.0158

Ceccarelli S, Grando S (1991) Environment of selection and type of germplasm in barley breeding for low-yielding conditions. Euphytica 57, 207–219. 10.1007/BF00039667

Ceccarelli S, Grando S, Baum M (2007) Participatory plant breeding in water-limited environments. Experimental Agriculture 43, 411–435. 10.1017/S0014479707005327

Challinor AJ, Watson J, Lobell DB, Howden SM, Smith DR, Chhetri N (2014) A meta-analysis of crop yield under climate change and adaptation. Nature Climate Change 4, 287–291. 10.1038/nclimate2153

Curti RN, Andrade AJ, Bramardi S, Velásquez B, Bertero DH (2012) Ecogeographic structure of phenotypic diversity in cultivated populations of quinoa from Northwest Argentina. Annals of Applied Biology 160, 114–125.

Curti RN, de la Vega AJ, Andrade AJ, Bramardi SJ, Bertero HD (2014) Multi- environmental evaluation for grain yield and its physiological determinants of quinoa genotypes across Northwest Argentina. Field Crops Research 166, 46–57. 10.1016/j.fcr.2014.06.011

Espitia-Rangel E (1994) Breeding of Grain Amaranth. In: Paredes-López O (ed) Amaranth Biology, Chemistry, and Technology, 1st edn. CRC Press Inc. Florida, pp 23-39

Ferrero ME, Villalba R (2019) Interannual and long-term precipitation variability along the subtropical mountains and adjacent Chaco (22–29° S) in Argentina. Frontiers in Earth Science 7. 10.3389/feart.2019.00148

Galluzzi G, López Noriega I (2014) Conservation and use of genetic resources of underutilized crops in the Americas—A continental analysis. Sustainability 6, 980–1017. 10.3390/su6020980

Gönen E, Çolak YB, Ozfidaner M, Yazar A, Tanriverdi Ç (2024) Eco-physiological response of grain amaranth to regulated deficit and conventional deficit irrigation applied with surface and subsurface drip irrigation systems. The Journal of Agricultural Science 1–15. 10.1017/S0021859624000480

Greizerstein EJ, Poggio L (1994) Karyological studies in grain *Amaranths*. Cytologia 59, 25–30. 10.1508/cytologia.59.25

Hatfield JL, Walthall CL (2015) Meeting global food needs: realizing the potential via genetics × environment × management interactions. Agronomy Journal 107, 1215–1226. 10.2134/agronj15.0076

Henderson TL, Johnson BL, Schneiter AA (1998) Grain amaranth seeding dates in the Northern Great Plains. Agronomy Journal 90, 339–344. 10.2134/agronj1998.00021962009000030005x

Hunziker AT (1952) Los pseudocereales de la agricultura indígena de América. Buenos Aires, Acme Jacobsen SE, Mujica, A., and Ortiz R (2003) The global potential for quinoa and other Andean crops. Food Reviews International 19, 139–148. 10.1081/FRI-120018880

Jamalluddin N, Massawe FJ, Mayes S, Ho WK, Singh A, Symonds RC (2021) Physiological screening for drought tolerance traits in vegetable amaranth (*Amaranthus tricolor*) germplasm. Agriculture 11, 994. 10.3390/agriculture11100994

Jamalluddin N, Massawe, Festo J, and Symonds RC (2019) Transpiration efficiency of Amaranth (*Amaranthus* sp.) in response to drought stress. The Journal of Horticultural Science and Biotechnology 94, 448–459. 10.1080/14620316.2018.1537725

Johnson BL, Henderson TL (2002) Water use patterns of grain amaranth in the Northern Great Plains. Agronomy Journal 94, 1437–1443. 10.2134/agronj2002.1437

Joshi DC, Sood S, Hosahatti R, Kant L, Pattanayak A, Kumar A, Yadav D, Stetter MG (2018) From zero to hero: the past, present and future of grain amaranth breeding. Theoretical and Applied Genetics 131, 1807–1823. 10.1007/s00122-018-3138-y

Kalinowski LS, Navarro JP, Concha AIR, Hermoza GC, Pacheco RA, Choquevilca YC, Jara EV (1992) Grain amaranth research in Peru. Food Reviews International 8, 87–124. 10.1080/87559129209540931

Liu F, Stützel H (2002) Leaf expansion, stomatal conductance, and transpiration of vegetable amaranth (*Amaranthus* sp.) in response to soil drying. Journal of the American Society for Horticultural Science 127, 878–883. 10.21273/JASHS.127.5.878

Liu F, Stützel H (2004) Biomass partitioning, specific leaf area, and water use efficiency of vegetable amaranth (*Amaranthus* spp.) in response to drought stress. Scientia Horticulturae 102, 15–27. 10.1016/j.scienta.2003.11.014

Lobell DB, Di Tommaso S (2025) A half-century of climate change in major agricultural regions: Trends, impacts, and surprises. Proceedings of the National Academy of Sciences 122, e2502789122. 10.1073/pnas.2502789122

Mujica A 1997 El cultivo del amaranto (*Amaranthus* spp.): producción, mejoramiento genetico y utilización, Universidad Nacional del Altiplano (UNA). Nieto C (1993) The preservation of foods indigenous to the Ecuadorian Andes. *Mountain Research and Development* **13**, 185–188. 10.2307/3673636

Olivoto T, Lúcio AD (2020) metan: An R package for multi-environment trial analysis. Methods in Ecology and Evolution 11, 783–789. 10.1111/2041-210X.13384

Olivoto T, Lúcio ADC, da Silva JAG, Marchioro VS, de Souza VQ, Jost E (2019) Mean performance and stability in multi-Environment trials I: combining features of AMMI and BLUP techniques. Agronomy Journal 111, 2949–2960. 10.2134/agronj2019.03.0220

Omami EN, Hammes PS, Robbertse PJ (2006) Differences in salinity tolerance for growth and water-use efficiency in some amaranth *(Amaranthus* spp.) genotypes. New Zealand Journal of Crop and Horticultural Science 34, 11–22. 10.1080/01140671.2006.9514382

Paderewski J, Gauch Jr. HG, Mądry W, Gacek E (2016) AMMI analysis of four-way genotype × location × management × year data from a wheat trial in Poland. Crop Science 56, 2157–2164. 10.2135/cropsci2015.03.0152

Padulosi S, Heywood V, Hunter D, Jarvis A (2011) Underutilized species and climate change: Current status and outlook. In ‘Crop Adaptation to Climate Change’. pp. 507–521. (John Wiley & Sons, Ltd)

Pulvento C, Lavini A, Riccardi M, d’Andria R, Ragab R (2015) Assessing amaranth adaptability in a Mediterranean area of South Italy under different climatic scenarios. Irrigation and Drainage 64, 50–58. 10.1002/ird.1906

Pulvento C, Sellami M houssemeddine, Lavini A (2022) Yield and quality of *Amaranthus hypochondriacus* grain amaranth under drought and salinity at various phenological stages in southern Italy. Journal of the Science of Food and Agriculture 102, 5022–5033. 10.1002/jsfa.11088

R Core Team. R: A Language and Environment for Statistical Computing. R Foundation for Statistical Computing, 2024. https://www.R-project.org/

Rojas W, Soto JL, Pinto M, et al (2010) Granos andinos: avances, logros y experiencias desarrolladas en quinua, cañahua y amaranto en Bolivia. Bioversity International, Roma, Italy. SAS (SAS Institute, Inc.). SAS/STAT® 9.4: User’s Guide, 2004.

Ulian T, Diazgranados M, Pironon S, Padulosi S, Liu U, Davies L, Howes M-JR, Borrell JS, Ondo I, Pérez-Escobar OA, Sharrock S, Ryan P, Hunter D, Lee MA, Barstow C, Łuczaj Ł, Pieroni A, Cámara-Leret R, Noorani A, Mba C, Nono Womdim R, Muminjanov H, Antonelli A, Pritchard HW, Mattana E (2020) Unlocking plant resources to support food security and promote sustainable agriculture. *PLANTS, PEOPLE*, PLANET 2, 421–445. 10.1002/ppp3.10145

Xangsayasane P, Jongdee B, Pantuwan G, Fukai S, Mitchell JH, Inthapanya P, Jothiyangkoon D (2014) Genotypic performance under intermittent and terminal drought screening in rainfed lowland rice. Field Crops Research 156, 281–292. 10.1016/j.fcr.2013.10.017

