## Supplementary material for "Harnessing drought tolerance in a reference set of Andean amaranths": Plot of performance for Andean amaranth genotypes evaluated in paired trials under irrigation and stressed conditions

**Figure S1** Plot of performance for Andean amaranth genotypes evaluated at four sites in Northwest Argentina in paired trials under irrigation and stressed conditions: (a) León; (b) Purmamarca; (c) Hornaditas; and (d) La Quiaca.

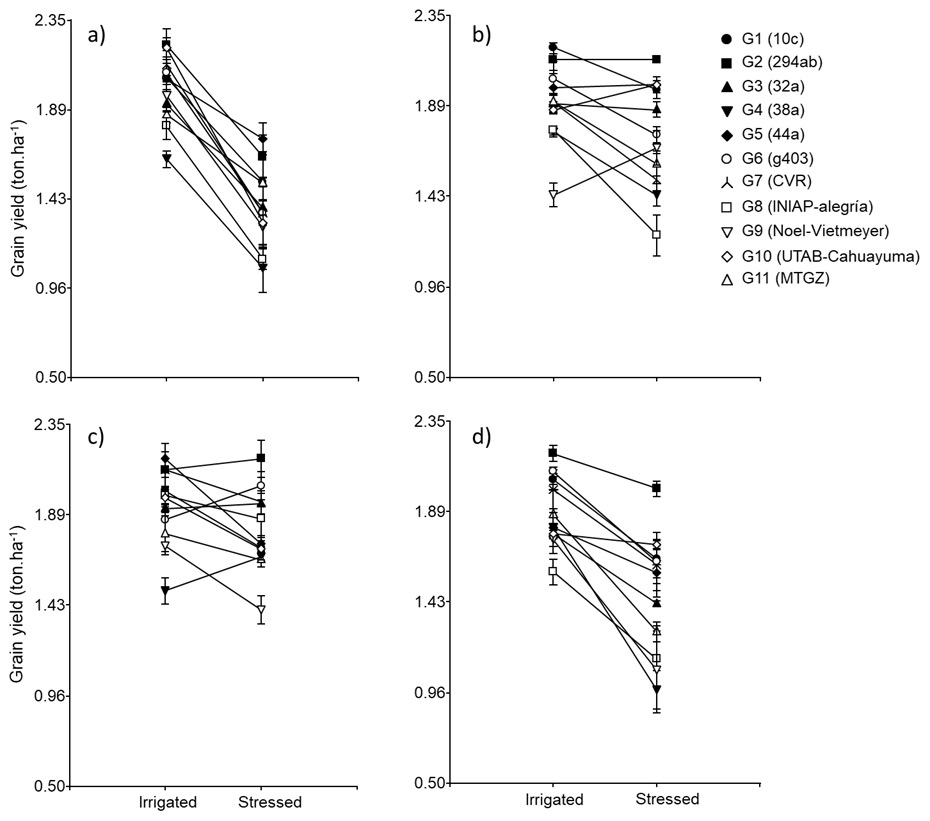
