## Supplementary material for "Harnessing drought tolerance in a reference set of Andean amaranths": The ANOVA for the additive main effects and multiplicative interaction model

**Table S1**. The ANOVA for the additive main effects and multiplicative interaction model

| Sources of variation | df | Sum of squares | Mean Squares | *F*-ratio | *P-*value | Proportion accumulated | |
| --- | --- | --- | --- | --- | --- | --- | --- |
| Environments (E) | 7 | 15.28 | 2.18 | 32.77 | 8.72e^-11^ |  |  |
| Genotypes (G) | 10 | 10.33 | 1.03 | 44.33 | 4.99e^-49^ |  |  |
| Block (E) | 24 | 1.60 | 0.07 | 2.86 | 2.37e^-05^ |  |  |
| G×E | 70 | 6.10 | 0.09 | 3.74 | 1.95e^-14^ |  |  |
| IPC1* | 16 | 2.37 | 0.15 | 6.35 | 0 | 38.8 | 38.8 |
| IPC2 | 14 | 1.21 | 0.09 | 3.70 | 0 | 19.8 | 58.6 |
| IPC3 | 12 | 1.16 | 0.10 | 4.16 | 0 | 19.1 | 77.7 |
| IPC4 | 10 | 0.65 | 0.07 | 2.80 | 2.70e^-03^ | 10.7 | 88.4 |
| IPC5 | 8 | 0.43 | 0.05 | 2.29 | 2.22e^-02^ | 7 | 95.4 |
| IPC6 | 6 | 0.24 | 0.04 | 1.69 | 1.24e^-01^ | 3.9 | 99.3 |
| IPC7 | 4 | 0.04 | 0.01 | 0.46 | 7.65e^-01^ | 0.7 | 100 |
| Residuals | 240 | 5.59 | 0.02 |  |  |  |  |

*Interaction Principal Component
